# Designing viscoelastic mucin-based hydrogels

**DOI:** 10.1101/656801

**Authors:** Katherine Joyner, Daniel Song, Robert F. Hawkins, Richard D. Silcott, Gregg A. Duncan

## Abstract

We report the design of a mucin hydrogel created using a thiol-based cross-linking strategy. By using a cross-linking reagent capable of forming disulfide linkages between mucins, the mucin-based hydrogels possess viscoelastic properties comparable to native mucus as measured by bulk rheology. We confirmed disulfide cross-links mediate gel formation in our system using chemical treatments to block and reduce cysteines where we found mucin hydrogel network formation was inhibited and disrupted, respectively. Particle tracking microrheology was used to investigate the kinetics and evolution of microstructure and viscoelasticity within the hydrogel as it formed. We found that the rate of gel formation could be tuned by varying the mucin to crosslinker ratio, producing network pore sizes in the range measured previously in human mucus. The results of this work provide a new, simple method for creating mucin hydrogels with physiologically relevant properties using readily available reagents.

## Introduction

Mucus is a biological gel that coats and protects epithelial surfaces in tissues throughout the body. Mucins are large, polymeric glycoproteins that are the primary contributor to the viscoelastic properties of mucus.^1-2^ The cysteine-rich domains of mucins facilitate their assembly into a mesh-like structure through disulfide cross-linking.^3-4^ This network architecture allows mucus to behave as a viscoelastic gel at the macroscale.^5-6^ On the microscopic scale, mucus acts as physical barrier, entrapping harmful pathogens and particulates to prevent them from reaching the underlying epithelium.^5-9^ However, the process by which mucins organize into a mucus gel has been challenging to recreate in the laboratory. As a result, our understanding of mucus properties is currently limited by the lack of suitable models. Given the natural anti-microbial properties of mucus,^10-12^ there has also been recent interest in engineering mucin-based materials for biomedical applications.^13-14^

In previous work, it has been observed that commercially available mucins (e.g. porcine gastric mucin, bovine submaxillary mucin) do not form hydrogels at physiological concentrations, presumably as a result of their processing.^15-16^ This considerably reduces their utility as a model of natural mucus. This has led to efforts to develop methods to purify mucins without compromising their capacity to form a gel.^17^ For example, it has been shown mucins can be purified from porcine stomach scrapings which preserves their natural gel-forming properties.^15^, ^18-21^ Mucus secreted from tissue culture models has also been collected and purified which has been shown to retain comparable viscoelastic behavior to human mucus.^22-23^ However, the requirement of specialized, relatively low-yield processing techniques limits their widespread usage. Methods to enable the formation of gels from crude, partially purified mucins available in large quantities from commercial vendors would allow for its general use for basic and applied research. In addition, it is challenging to control the degree of disulfide cross-linking within these gels which will directly influence their functional properties.

Towards this end, we have developed a simple approach to generate hydrogels using crude, commercially available mucins and thiol-based cross-linkers that re-establish their native rheological properties. Particle tracking microrheology (PTM) of PEG-coated nanoparticles was used to determine if and when mucins began to form into a gel after addition of a cross-linking reagent. Several thiol-functionalized, polymer-based crosslinkers with varying chemistry and geometry were tested to determine which would initiate mucin hydrogel network formation. Using an optimized cross-linking strategy, bulk rheological assessment of mucin-based hydrogels was performed to determine if we were able to produce gels with physiological viscoelastic properties. Finally, PTM was used to investigate the kinetics of gel formation as a function of mucin and cross-linker concentration. The results of this work demonstrate how these mucins, incapable of forming viscoelastic gels, may be restored to behave like natural mucus.

## Experimental

### Nanoparticle preparation

Carboxylate modified fluorescent PS spheres (PS-COOH; Life Technologies) with a diameter of 100 or 500 nm were coated with a high surface density of polyethylene glycol (PEG) via a carboxyl-amine linkage using 5-kDa methoxy PEG-amine (Creative PEGWorks) as previously reported.^8^, ^24^ Briefly, PEG-amine was added to a diluted suspension of PS-COOH in ultrapure water. N-hydroxysulfosucciniminde sodium salt (10 mM; Alfa Aesar), borate buffer (pH 8.3) and 1-ethyl-3-(3-dimethylaminopropyl) carbodiimide hydrochloride (2 mM; Thermo Fisher) were then added to activate and link PEG-amine to PS-COOH nanoparticles. The reaction was mixed for at least 4 hours at room temperature and covered to minimize exposure to light. After mixing, excess reagents were removed by washing and centrifuging three times with ultrapure water, then re-suspended to a final volume two times the original. After washing, particle size and zeta potential was measured in 10 mM NaCl at pH 7 using a Malvern Zetasizer Nano ZS90. We measured diameters of 122 nm and 428 nm and zeta potentials of -4.22 and -3.4 mV for 100 nm and 500 nm PEG-coated PS nanoparticles, respectively.

### Mucin-based hydrogel preparation

Solutions of porcine gastric mucin (PGM; Sigma-Aldrich) or mucin from bovine submaxillary glands (BSM; Sigma-Aldrich) were prepared in a physiological buffer (154 mM NaCl, 3 mM CaCl_2_, and 15 mM NaH_2_PO_4_ at pH 7.4) and stirred at room temperature for 2 hours. Crosslinking reagents tested in this work include monofunctional PEG-thiol (PEG-1SH; 10 kDa), bifunctional PEG-thiol (PEG-2SH; 10 kDa), 4-arm PEG-thiol (PEG-4SH; 10 kDa), each purchased from Laysan Bio Inc. and a thiol-functionalized dextran (dextran-SH; 10 kDa) purchased from Nanocs. Cross-linking solutions were prepared separately in buffer directly before mixing with mucin solutions. To initiate gelation, equal volume aliquots of each solution were mixed and either equilibrated for 30 minutes before particle tracking or left for 24 hours before bulk rheological characterization at room temperature. To confirm cross-linker interaction with cysteine rich regions of PGM, thiol-blocking and disulfide cleavage experiments were performed to inhibit or disrupt gel assembly. Iodoacetamide at 50 mM (VWR) was added to the PGM solutions and mixed for 1 hour to alkylate PGM cysteine residues before mixing with the crosslinking polymer.^25^ As has been previously shown in human mucus,^26^ we disrupted disulfide bonds within the gel using the reducing agent, *N*-acetyl cysteine (NAC). NAC was added after gelation (>9 hours) at a final concentration of 50 mM (NAC) and incubated at 37°C for 30 minutes.

### Bulk rheological measurements

Dynamic rheological measurements of the mucin/PEG-4SH hydrogels were performed using an AR2000 rheometer (TA Instruments) with a 20-mm diameter parallel plate geometry at 25°C. To determine the linear viscoelastic region of the fully formed gel, a strain sweep measurement was collected from 0.1-10% strain at a frequency of 1 rad/s. To determine the elastic modulus, *G*’(ω), and viscous modulus, *G*”(ω), a frequency sweep measurement was conducted within the linear viscoelastic region of the gel, at 1% strain amplitude and angular frequencies from 0.1 to 100 rad/s.

### Sample preparation for fluorescent video microscopy

The diffusion of PEG-coated nanoparticles (PEG-NP) in mucin-based hydrogels were measured using fluorescent video microscopy. Samples were prepared with 0.5 μl of ∼0.002% w/v suspension of PEG-NP, added into a 25 µL solution of PGM or BSM and cross-linker prior to gel formation. Next, ∼25µL of mucin hydrogel precursor solution containing PEG-NP was added into a custom microscopy chamber, sealed with a cover slip, and equilibrated for 30 minutes at room temperature before imaging. Fluorescent video microscopy experiments were collected using a Zeiss 800 LSM microscope equipped with a ×63/1.20 W Korr UV VIS IR water-immersion objective and an Axiocam 702 camera with pixel resolution of 0.093µm/pixel. Images were collected at a frame rate of 33.33 Hz for 10 seconds at room temperature. For studies of gel kinetics, videos were taken every hour in order to measure PEG-NP diffusion during hydrogel assembly.

### Particle tracking microrheology (PTM)

The particle tracking data analysis used in this work is based on a previously developed image processing algorithm.^27^ To briefly describe, an image intensity threshold is set to differentiate nanoparticle signal from background. Nanoparticles above the intensity threshold are then assigned centers based on *x* and *y* image location. Particles were tracked for at least 300 sequential frames allowing trajectories to be determined. From these trajectories, time-averaged mean squared displacement (MSD; ⟨Δ*r*^2^(τ)⟩) as a function of lag time, τ, was calculated as ⟨Δ*r*^2^(*τ*)⟩ = ⟨[*x*(*t*+*τ*)-*x*(*t*)]^2^+[*y*(*t+τ*)-*y*(*t*)]^2^⟩.

Using the generalized Stokes-Einstein relation,^28^ measured MSD values were used to compute viscoelastic properties of the hydrogels. The Laplace transform of ⟨Δ*r*^2^(τ)⟩, ⟨Δ*r*^2^(s)⟩, is related to viscoelastic spectrum 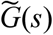 using the equation 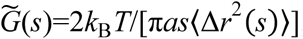, where *k*_B_*T* is the thermal energy, *a* is the particle radius, *s* is the complex Laplace frequency. The complex modulus can be calculated as *G*(*ω)=*G*’(ω)+*G*”*(i*ω), with iω being substituted for *s*, where *i* is a complex number and ω is frequency. Hydrogel network pore size, ξ, is estimated based on MSD using the equation, 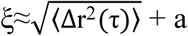,^29^ and also based on *G*’ using the equation, ξ≈(*k*_B_*T*/G’)^1^/^3^.^30^ The sol-gel transition (gel point) was determined by calculating the logarithmic slope of the mean-squared displacement, α=log_10_(⟨Δr^2^(τ)⟩)/log_10_(τ). We used a critical value of *n*=0.5 where the gel point is defined as the time when measured α < *n*.^31^

## Results & Discussion

### Design and characterization of mucin-based hydrogel network assembly

We aimed to create a viscoelastic gel composed of porcine gastric mucin (PGM) via disulfide-bond mediated crosslinking between the cysteine domains of PGM biopolymers. PTM was employed to evaluate different strategies to make mucin-based gels. We chose to use PTM as it provides a biophysical method to assess the mechanical properties of soft biological gels. Importantly, it has been used extensively in previous work to evaluate microrheology and structure of mucus.^19^, ^22^, ^24^, ^26^, ^32-34^ The diffusion of fluorescently labeled, PEG-coated particles (PEG-NP) with diameter of 100 nm was measured in mixtures of PGM and various thiol-containing polymers to determine which may be a suitable cross-linking reagent. We used a low concentration (∼4 × 10^-5^ % w/v) of nanoparticles coated with a dense layer of low molecular weight PEG in order to ensure nanoparticle probes are minimally adhesive to mucins.^35-37^ Thus, PEG-NP probes are unlikely to interfere or influence hydrogel formation. Figure 1 shows PEG-NP diffusion as measured by the mean squared displacement (MSD; ⟨Δ*r*^2^⟩) of the particles determined immediately after mixing (hour 0) and 9 hours after mixing. Once the gel has formed, the MSD of PEG-NP will be reduced due to steric interactions with the mesh-like mucin polymer network. In addition, the logarithmic slope of MSD will be reduced from ∼1 for a viscous liquid to ≤ 0.5 for a viscoelastic material.^33^, ^38^

**Figure 1.**
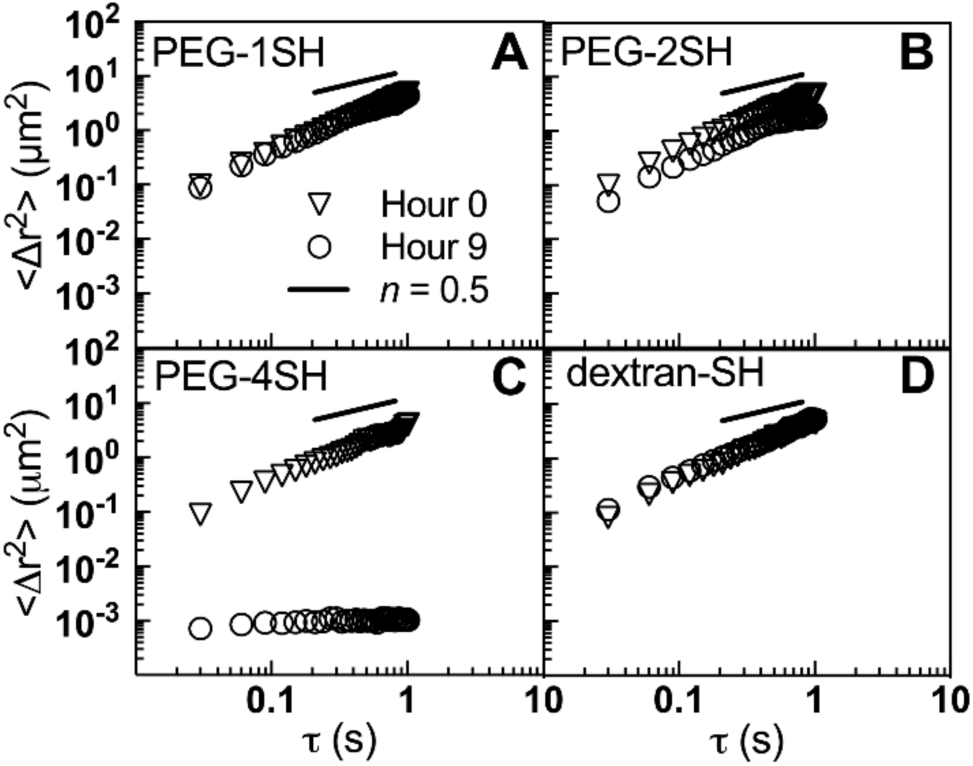
Microrheology of mucin mixed with various thiol-based crosslinkers. **(A-D)** Multiple particle tracking of 100-nm PEG-NP in mucin solutions with varying cross-linking polymers. Each panel represents the ensemble average mean squared displacement as function of time scale τ (MSD; ⟨Δ*r*^2^⟩) of 100 nm PEG-NP at hour 0 (triangles) and hour 9 (circles) in 2% w/v of PGM and 2% w/v of (**A**) PEG-1SH, (**B**) PEG-2SH, (**C**) PEG-4SH, and (**D**) dextran-SH. Reference lines with slope *n*=0.5 are included in each panel.

Four potential thiol-based crosslinking reagents were tested with a low molecular weight of 10 kDa and a PEG or dextran backbone. Two percent w/v of mucin and crosslinker was used in each case. As shown in Fig. 1A, no significant changes in MSD of PEG-NP were observed after mixing PGM with monofunctional PEG-1SH. We hypothesized bifunctional PEG-2SH would be able to create disulfide bridges between mucin polymers to facilitate gel formation. However, we found it was unable to form a gel over a 9-hr period (Fig. 1B). Using a multi-arm PEG-4SH, we observed a substantial decrease in the magnitude and slope of MSD after 9 hours of incubation with PGM, indicative of network formation (Fig. 1C). We also considered the potential self-assembly of PEG-4SH, yielding a hydrogel that does not incorporate mucin biopolymers. To address this, we first measured the microrheology of a 2% PEG-4SH solution (i.e. in the absence of PGM), where no gel formation was observed in a 9-hour period (Fig. S1). We also examined if PEG-4SH self-assembly could be the result of an increased local concentration due to crowding by mucin polymers. To evaluate this, 500 kDa hyaluronic acid (HA) was used to act as a non-thiol containing, high molecular weight crowding agent. Hydrogel network formation was not observed after a 9-hour period in mixtures of 2% HA and 2% PEG-4SH (Fig. S2), suggesting crowding did not induce PEG-4SH self-assembly. We then changed the cross-linking polymer backbone from PEG to dextran, but unlike PEG-4SH, the multi-functional dextran-SH did not lead to assembly of mucins into a mesh network (Fig. 1D). These results suggest that the more flexible PEG backbone may also be important in the ability of PEG-4SH to facilitate hydrogel formation. Based on these results, PEG-4SH was selected as the optimal cross-linking polymer to produce mucin-based hydrogels.

In order to examine the mechanism of gel formation, we tested if alkylation or reduction of cysteines impacted the formation of mucin-based hydrogels. PTM was again employed to monitor changes in the mucin hydrogel network upon treatment with cysteine-blocking or disulfide bond-disrupting agents. In order to block available cysteines, a 2% PGM solution was alkylated with iodoacetamide (IAM). While only the PGM solution was treated with IAM, we note excess IAM would most likely block cysteines present on PEG-4SH once the solutions were mixed. As shown in Fig. 2A, no significant changes in 100 nm PEG-NP diffusion were observed after 9 hours, implying network assembly is inhibited by IAM treatment. To determine if the hydrogel network would be disrupted by cleaving disulfide bonds, the reducing reagent, *N*-acetyl cysteine (NAC), was applied to a fully formed PGM/PEG-4SH hydrogel. After 30-minute treatment with NAC, we observed a marked increase in 100 nm PEG-NP diffusion approaching that of the PGM solution alone (Fig. 2B), indicative of a significant breakdown in the mesh network. Together, these data further supports that disulfide bonds between PGM and PEG-4SH drive network assembly.

**Figure 2.**
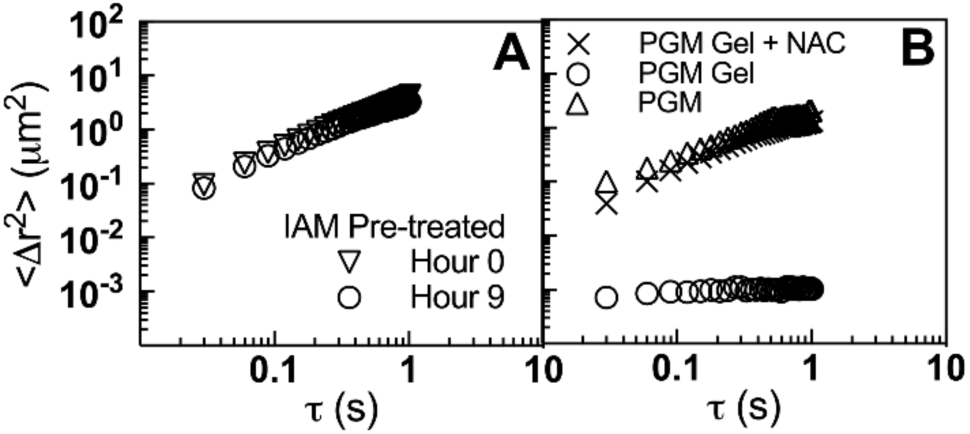
Effects of cysteine-blocking and disulfide cleavage on mucin-based hydrogel assembly and disassembly. (**A**) MSD (⟨Δ*r*^2^⟩) as a function of time scale (τ) of 100 nm PEG-NP in 2% w/v PGM and 2% w/v PEG-SH-4 solution 0 hours (downward triangles) and 9 hours (circles) after pre-treatment with iodoacetamide (IAM). (**B**) MSD as a function of τ of 100 nm PEG-NP in 2% w/v PGM solution after 9 hours (PGM; upward triangles), in 2% PGM/PEG-4SH hydrogel after 9 hours (PGM Gel; circles), and in 2% PGM/PEG-4SH hydrogel after a 30-minute treatment with *N*-acetyl cysteine (PGM Gel + NAC; cross).

### Macro- and microrheology of mucin-based hydrogels

To determine if a viscoelastic gel similar to native mucus was formed, we measured the bulk elastic (*G*’) and viscous (*G*”) moduli of 2% PGM after mixing and incubation for 24 hours with 2% PEG- 4SH. Strain sweep experiments were performed at frequency of ω=1 rad/s from 0.1 to 10% strain as shown in Fig. 3A. Elastic solid-like behavior of the material was observed with *G’* > *G”* and loss tangent (tan δ) of 8.1, thus confirming the viscoelastic nature of the hydrogel. We did not find evidence of yielding and thus, were within the linear viscoelastic region of the gel. To determine if the behavior of our hydrogel was frequency dependent, a frequency sweep was performed at 1% strain over a physiologically relevant frequency range of 0.1 – 100 rad/s. We confirmed the hydrogel was stable over the full range of frequencies tested (Fig. 3B). Importantly, we found the elastic modulus of ∼244 Pa at ω=1 rad/s is within the physiological range for mucus.^5^, ^24^, ^33^ These results suggest that PEG-4SH is able to support gelation of mucins into a viscoelastic network. To determine if this approach was generalizable to other mucin types, we prepared a hydrogel using another commercially available mucin, bovine submaxillary mucin (BSM). We measured the bulk elastic (*G*’) and viscous (*G*”) moduli of 2% BSM after mixing and incubation for 24 hours with 2% PEG-4SH. We also observed formation of a stable viscoelastic hydrogel with *G’* > *G”* and loss tangent (tan δ) of 1.4 (Fig. 3C,D).

**Figure 3.**
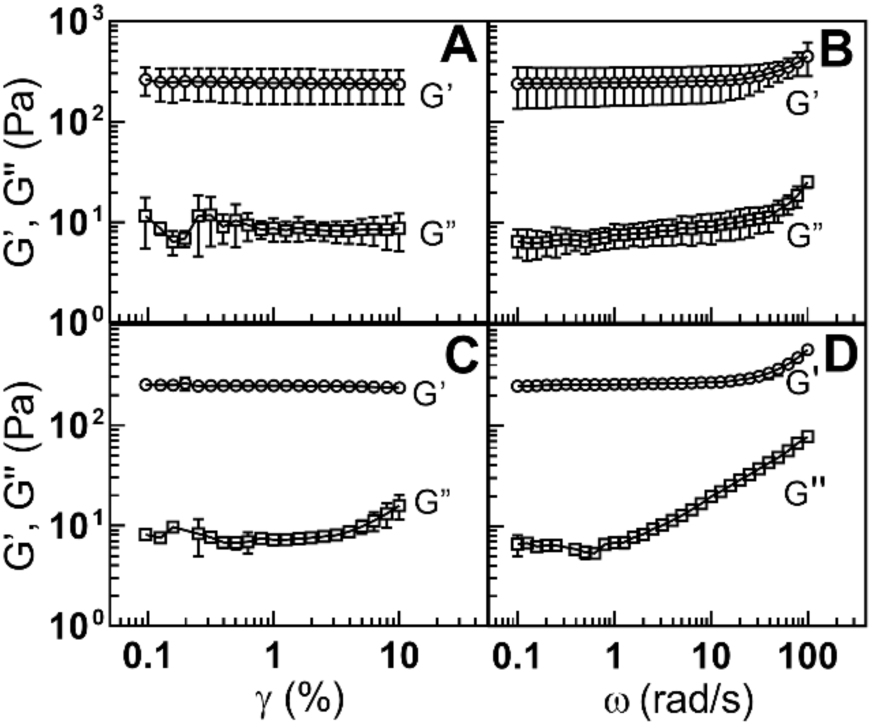
Bulk rheological characterization of mucin-based hydrogels. Elastic modulus *G’* (circles) and viscous modulus, *G”* (squares) of PGM-based **(A**,**B)** and BSM-based **(C**,**D)** mucin hydrogels after a 24-hr gelation time. Data represents average of 3 independently prepared hydrogels. (**A**) Strain (γ) sweep at frequency (ω) of 1 rad/s and (**B**) frequency sweep at γ=1% for 2% PGM and 2% PEG-4SH hydrogel. At ω = 1 rad/s, we measured *G*’ ∼ 244 Pa and tan δ ∼ 8.1. (**C**) Strain (γ) sweep at frequency (ω) of 1 rad/s and (**D**) Frequency sweep at γ=1% for 2% BSM and 2% PEG-4SH hydrogel. At ω = 1 rad/s, we measured *G*’ ∼ 270 Pa and tan δ ∼ 1.4.

Next, PTM was used to gain additional insights into the biophysical properties of the material. Given previous reports showing native mucus having network pore sizes ranging from 100-500 nm,^8^, ^34^, ^39^ we compared microrheology of a 2% PGM and 2% PEG-4SH hydrogel as measured using PEG-NP probes with 100 and 500 nm diameter. PTM measurements were performed each hour for a total of 9 hours to capture local changes in viscosity caused by the organization of mucin fibers into a hydrogel network over time. Figures 4A and 4B show MSD at τ = 1 s for 100 nm and 500 nm PEG-NP, respectively. A decrease in MSD as a function of gelation time was observed for both the 100 and 500 nm PEG-NP, indicating a decrease in diffusion rate as a result of gel network formation. Next, the ratio of elastic and viscous moduli (*G*’/*G*”) was calculated based on 100 nm (Fig. 4C) and 500 nm (Fig. 4D) PEG-NP probes. It should be noted these local measurements of *G*’ and *G*” will strongly depend on probe size as the gel network evolves over time. Based on measured MSD and *G*’, approximate hydrogel network mesh pore size were found to be on the order of microns at hour 0 and steadily decreasing with median pore sizes on the order of 200-300 nm, consistent with findings in mucus collected from humans.^8^, ^34^, ^39-40^

**Figure 4.**
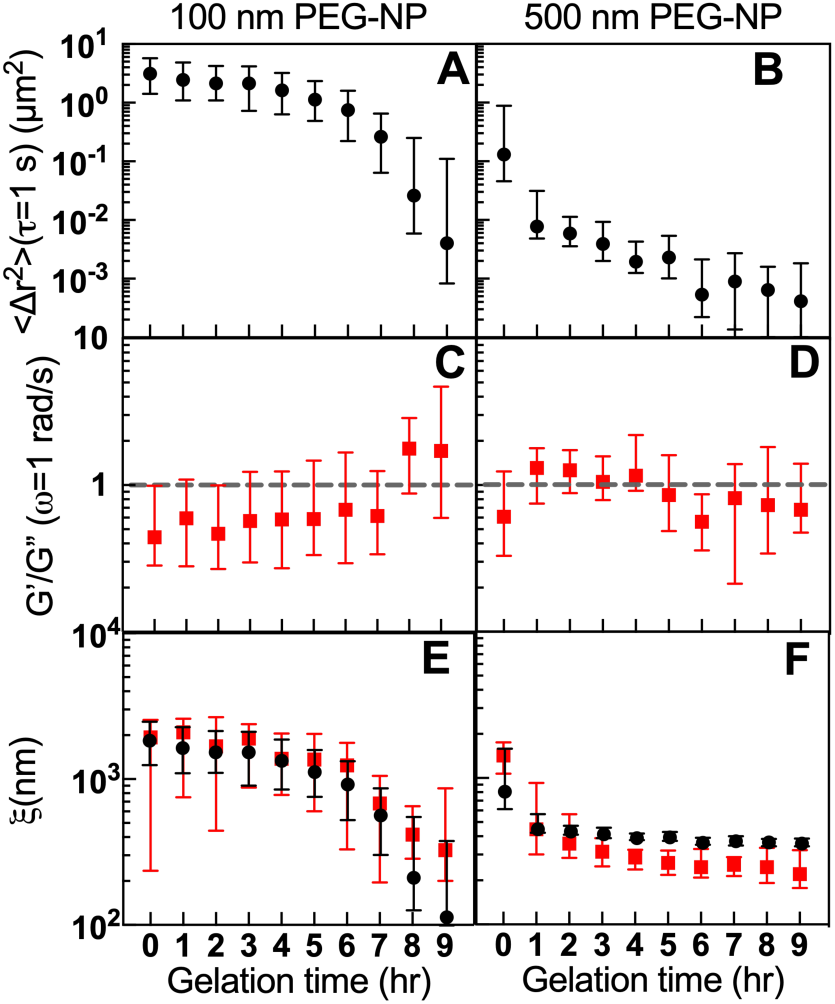
Impact of probe size on the microrheology of mucin-based hydrogels. **(A-F)** Microrheology of a 2% w/v PGM and 2% PEG-4SH gel measured using PEG-NP probes with diameters of 100 and 500 nm. Symbols in each panel are median values and bars indicate the interquartile range. (**A**,**B**) The MSD at τ = 1 s (⟨Δ*r*^2^(τ=1 s)⟩) for **(A)** 100 and **(B)** 500 nm PEG-NP during a 9-hour gelation period. (**C**,**D**) Elastic to viscous moduli ratio (G’/G”) at ω = 1 rad/sec calculated from measured MSD of **(C)** 100 nm and **(D)** 500 nm PEG-NP.(**E**,**F**) Estimated pore size based on analysis of MSD at 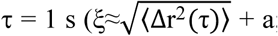; black circles) and G’ at ω = 1 rad/s (ξ≈(*k*B*T*/G’)^1^/^3^; red squares) for **(E)** 100 nm and **(F)** 500 nm PEG-NP probes.

Using nanoparticle probes of different size enabled us to resolve the early and late stages of mucin hydrogel assembly. Based on this analysis, we find evidence of distinct phases in mucin organization into a hydrogel network (i.e. transitions in *G*’/*G*” to values > 1). After ∼1 hour, 500 nm probes detect an initial rise in elasticity and reduction of network pore sizes (Fig. 4D,F). Starting at approximately hour 7, further reductions in network pore size are evident based on the relative increase in *G*’/*G*” measured by 100 nm probes (Fig. 4C, E). These data may indicate the mucin hydrogel forms with a rate of gelation that slows as network pore dimensions decrease. In future work, microrheological analysis of gel kinetics with greater resolution in time (i.e. measurements taken on the order of minutes) will be used to further interrogate mucin gel assembly. In the next section, we will further discuss gelation kinetics and how we may manipulate the sol-gel transition of these hydrogels.

### Concentration dependence of the sol-gel transition

To understand the influence of mucin to crosslinker ratio on gel assembly, PGM and PEG-4SH concentration were systematically varied to assess their impact on gelation kinetics. The time of gelation was determined based on measured MSD of 100 nm PEG-NP defined as α= log_10_[MSD]/log_10_[τ] = 0.5. Given 100 nm PEG-NP captured the latter stages of gel assembly (Fig. 4), probes of this size were used to monitor changes in α every hour for 9 hours as the network assembled. PGM concentration was varied 1-5% w/v while keeping the PEG-4SH concentration constant at 2% w/v. The data in Fig. 5A shows that increasing PGM concentration leads to increases in the rate of gelation evident by measured α values. Next, PEG-4SH concentration was varied from 1-5% w/v, while keeping PGM concentration constant at 2% w/v. We also observed a concentration dependent change in gelation kinetics with a reduction in the time needed to form a gel as PEG-4SH concentration is increased (Fig. 5B). Each series of results with data from individual experiments are provided in the supplemental figures (Fig. S3,4).

**Figure 5.**
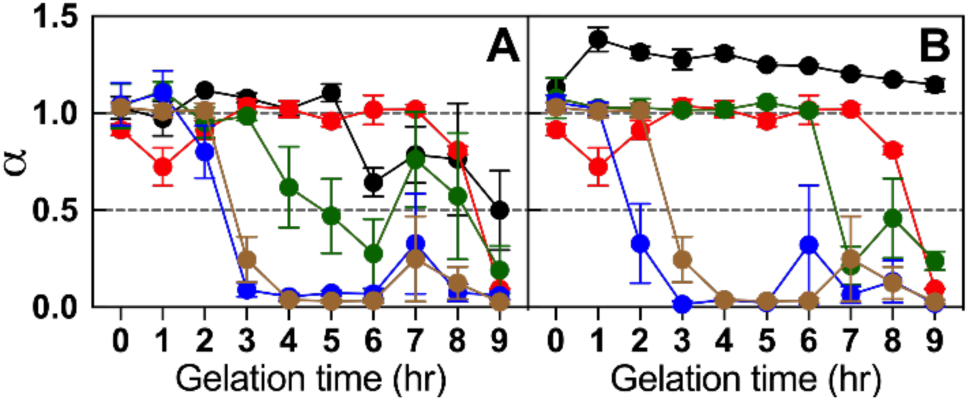
Gelation rate of mucin-based hydrogels with varying PGM and PEG-4SH concentration. **(A,B)** Kinetics of PGM/PEG-4SH gel formation as analyzed by microrheology using 100 nm PEG-NP probes. The gelation point is measured as α=log_10_[MSD]/log_10_[τ] ≤ 0.5. Each panel displays the mean and standard error of measured α. **(A)** α as a function of gelation time in hydrogels with a constant PEG-4SH concentration of 2% w/v and varying PGM concentrations of 1% w/v (black), 2% w/v (red), 3% w/v (green), 4% w/v (blue), and 5% w/v (brown). **(B)** α as a function of gelation time in hydrogels with a constant PGM concentration of 2% w/v and varying PEG-4SH concentration of 1% w/v (black), 2% w/v (red), 3% w/v (green), 4% w/v (blue), and 5% w/v (brown).

These results are consistent with our earlier findings, showing the gelation process is dependent on both PGM and PEG-4SH to create the viscoelastic gel. As one would intuitively expect, increasing the concentration of either mucin or crosslinking polymers leads to a decrease in time to form the gel. Starting from either 1% PGM or 1% PEG-4SH, we appear to be below the threshold concentration required to form a gel within a 9-hour period. However, we note the 1% PGM does approach the gel point near hour 9. As PGM concentration increases, a steady decrease in gelation time going is observed from an initial gel point of ∼9 hours for 2% w/v PGM, to ∼5 hours for 3% w/v PGM and to ∼3 hours for 5% w/v PGM (Fig. 5A). This can be simply explained as PGM concentration increases, the closer proximity between fibers makes PEG-4SH-mediated bridging more probable and as a result, gel assembly is more rapid. Interestingly, a more step-wise response in gelation time is observed as a function of PEG-4SH concentration with ∼8-9 hour for the lower 2% and 3% PEG-4SH concentrations and much faster ∼2-3 hr gelation times for the higher 4% and 5% PEG-4SH concentrations (Fig. 5B). This might be explained as PEG-4SH attach to the mucin fibers at higher density, the rate of assembly is enhanced due to an increase in their multi-valent bridging capacity.

Our results demonstrate how the mucin-based hydrogel may be tuned to assemble in a user-defined time frame. This presents the opportunity for our hydrogel system to be designed for various biomedical applications where slower kinetics (e.g. cell encapsulation^41^) or more rapid kinetics (e.g. injectable gels^42^) may be desirable. This system may also be used for fundamental studies on the role of kinetics in mucus gel formation and its biological function, which due to the lack of available models has not been previously investigated. While the results here motivate the continued optimization of the system, there are limitations in the present work which should be addressed in the future. First, we have yet to explore the effects of buffer conditions (e.g. ionic strength, pH) which can vary significantly between mucosal tissues and it will be important to evaluate this in our future work. Second, we have yet to determine if our synthetic system functionally behaves like mucus in the body. For example, we plan to investigate properties such as tissue adhesion, pathogen capture, and transportability of the gel in response to shear as these are all important aspects of mucus function.

## Conclusions

A mucin-based synthetic hydrogel was designed and characterized using particle tracking microrheology. We determined that hydrogel assembly was mediated by disulfide linkages between porcine gastric mucin and 4-arm PEG-thiol polymers. Bulk rheology confirmed the formation of a solid-like viscoelastic material, with an elastic modulus resembling naturally produced mucus. During the assembly process, the evolution of microscale structure in the mucin gel network was apparent based on microrheological measurements using PEG-coated nanoparticle probes of different sizes. We also found gelation kinetics can be tailored by changing mucin or cross-linker concentrations. In future work, we hope to further investigate how changes in gelation kinetics may influence mucin fiber structure and network heterogeneity. The results in this work establishes a simple means to produce mucin-based hydrogels which may be useful for a range of biological and medical applications.

## Acknowledgements

This work was supported by a Burroughs Wellcome Fund Career Award at the Scientific Interface (to GAD) and the University of Maryland.

## References

1. Bansil, R.; Turner, B. S., The biology of mucus: Composition, synthesis and organization. Advanced drug delivery reviews 2018, 124, 3–15.

2. Linden, S.; Sutton, P.; Karlsson, N.; Korolik, V.; McGuckin, M., Mucins in the mucosal barrier to infection. Mucosal immunology 2008, 1 (3), 183.

3. Thornton, D. J.; Sheehan, J. K., From mucins to mucus: toward a more coherent understanding of this essential barrier. Proceedings of the American Thoracic Society 2004, 1 (1), 54–61.

4. Perez-Vilar, J.; Hill, R. L., The structure and assembly of secreted mucins. Journal of Biological Chemistry 1999, 274 (45), 31751–31754.

5. Lai, S. K.; Wang, Y.-Y.; Wirtz, D.; Hanes, J., Micro-and macrorheology of mucus. Advanced drug delivery reviews 2009, 61 (2), 86–100.

6. Cone, R. A., Barrier properties of mucus. Advanced drug delivery reviews 2009, 61 (2), 75–85.

7. Carlson, T.; Lock, J.; Carrier, R., Engineering the mucus barrier. Annual review of biomedical engineering 2018, 20, 197–220.

8. Duncan, G. A.; Jung, J.; Hanes, J.; Suk, J. S., The mucus barrier to inhaled gene therapy. Molecular Therapy 2016, 24 (12), 2043–2053.

9. Berry, M.; Corfield, A. P.; McMaster, T. J., Mucins: a dynamic biology. Soft Matter 2013, 9 (6), 1740–1743.

10. Lieleg, O.; Lieleg, C.; Bloom, J.; Buck, C. B.; Ribbeck, K., Mucin biopolymers as broad-spectrum antiviral agents. Biomacromolecules 2012, 13 (6), 1724–1732.

11. Co, J. Y.; Cárcamo-Oyarce, G.; Billings, N.; Wheeler, K. M.; Grindy, S. C.; Holten-Andersen, N.; Ribbeck, K., Mucins trigger dispersal of Pseudomonas aeruginosa biofilms. NPJ biofilms and microbiomes 2018, 4 (1), 23.

12. Caldara, M.; Friedlander, R. S.; Kavanaugh, N. L.; Aizenberg, J.; Foster, K. R.; Ribbeck, K., Mucin biopolymers prevent bacterial aggregation by retaining cells in the free-swimming state. Current Biology 2012, 22 (24), 2325–2330.

13. Duffy, C. V.; David, L.; Crouzier, T., Covalently-crosslinked mucin biopolymer hydrogels for sustained drug delivery. Acta biomaterialia 2015, 20, 51–59.

14. Yan, H.; Chircov, C.; Zhong, X.; Winkeljann, B.; Dobryden, I.; Nilsson, H. E.; Lieleg, O.; Claesson, P. M.; Hedberg, Y.; Crouzier, T., Reversible condensation of mucins into nanoparticles. Langmuir 2018, 34 (45), 13615–13625.

15. Celli, J. P.; Turner, B. S.; Afdhal, N. H.; Ewoldt, R. H.; McKinley, G. H.; Bansil, R.; Erramilli, S., Rheology of gastric mucin exhibits a pH-dependent solgel transition. Biomacromolecules 2007, 8 (5), 1580–1586.

16. Kočevar-Nared, J.; Kristl, J.; Šmid-Korbar, J., Comparative rheological investigation of crude gastric mucin and natural gastric mucus. Biomaterials 1997, 18 (9), 677–681.

17. Davies, J. R.; Carlstedt, I., Isolation of large gel-forming mucins. In Glycoprotein Methods and Protocols, Springer: 2000; pp 3–13.

18. Schömig, V. J.; Käsdorf, B. T.; Scholz, C.; Bidmon, K.; Lieleg, O.; Berensmeier, S., An optimized purification process for porcine gastric mucin with preservation of its native functional properties. RSC Advances 2016, 6 (50), 44932–44943.

19. Wagner, C. E.; Turner, B. S.; Rubinstein, M.; McKinley, G. H.; Ribbeck, K., A rheological study of the association and dynamics of MUC5AC gels. Biomacromolecules 2017, 18 (11), 3654–3664.

20. Cao, X.; Bansil, R.; Bhaskar, K. R.; Turner, B. S.; LaMont, J. T.; Niu, N.; Afdhal, N. H., pH-dependent conformational change of gastric mucin leads to sol-gel transition. Biophysical journal 1999, 76 (3), 1250–1258.

21. Georgiades, P.; Pudney, P. D.; Thornton, D. J.; Waigh, T. A., Particle tracking microrheology of purified gastrointestinal mucins. Biopolymers 2014, 101 (4), 366–377.

22. Hill, D. B.; Vasquez, P. a.; Mellnik, J.; McKinley, S. a.; Vose, A.; Mu, F.; Henderson, A. G.; Donaldson, S. H.; Alexis, N. E.; Boucher, R. C.; Forest, M. G., A biophysical basis for mucus solids concentration as a candidate biomarker for airways disease. PLoS ONE 2014, 9, 1–11.

23. Button, B.; Goodell, H. P.; Atieh, E.; Chen, Y.-C.; Williams, R.; Shenoy, S.; Lackey, E.; Shenkute, N. T.; Cai, L.-H.; Dennis, R. G., Roles of mucus adhesion and cohesion in cough clearance. Proceedings of the National Academy of Sciences 2018, 115 (49), 12501–12506.

24. Schuster, B. S.; Suk, J. S.; Woodworth, G. F.; Hanes, J., Nanoparticle diffusion in respiratory mucus from humans without lung disease. Biomaterials 2013, 34 (13), 3439–46.

25. Sechi, S.; Chait, B. T., Modification of cysteine residues by alkylation. A tool in peptide mapping and protein identification. Analytical Chemistry 1998, 70 (24), 5150–5158.

26. Suk, J. S.; Lai, S. K.; Boylan, N. J.; Dawson, M. R.; Boyle, M. P.; Hanes, J., Rapid transport of muco-inert nanoparticles in cystic fibrosis sputum treated with N-acetyl cysteine. Nanomedicine 2011, 6 (2), 365–375.

27. Schuster, B. S.; Ensign, L. M.; Allan, D. B.; Suk, J. S.; Hanes, J., Particle tracking in drug and gene delivery research: State-of-the-art applications and methods. Adv Drug Deliv Rev 2015, 91, 70–91.

28. Mason, T. G.; Weitz, D., Optical measurements of frequency-dependent linear viscoelastic moduli of complex fluids. Physical review letters 1995, 74 (7), 1250.

29. Shin, J. H.; Gardel, M.; Mahadevan, L.; Matsudaira, P.; Weitz, D., Relating microstructure to rheology of a bundled and cross-linked F-actin network in vitro. Proceedings of the National Academy of Sciences 2004, 101 (26), 9636–9641.

30. De Gennes, P.-G.; Gennes, P.-G., Scaling concepts in polymer physics. Cornell university press: 1979.

31. Savin, T.; Doyle, P. S., Electrostatically tuned rate of peptide self-assembly resolved by multiple particle tracking. Soft Matter 2007, 3 (9), 1194–1202.

32. Duncan, G. A.; Jung, J.; Joseph, A.; Thaxton, A. L.; West, N. E.; Boyle, M. P.; Hanes, J.; Suk, J. S., Microstructural alterations of sputum in cystic fibrosis lung disease. JCI Insight 2016, 1 (18), e88198.

33. Dawson, M.; Wirtz, D.; Hanes, J., Enhanced viscoelasticity of human cystic fibrotic sputum correlates with increasing microheterogeneity in particle transport. The Journal of biological chemistry 2003, 278, 50393–401.

34. Lai, S. K.; Wang, Y. Y.; Hida, K.; Cone, R.; Hanes, J., Nanoparticles reveal that human cervicovaginal mucus is riddled with pores larger than viruses. Proc Natl Acad Sci U S A 2010, 107 (2), 598–603.

35. Xu, Q.; Ensign, L. M.; Boylan, N. J.; Schön, A.; Gong, X.; Yang, J.-C.; Lamb, N. W.; Cai, S.; Yu, T.; Freire, E., Impact of surface polyethylene glycol (PEG) density on biodegradable nanoparticle transport in mucus ex vivo and distribution in vivo. ACS nano 2015, 9 (9), 9217–9227.

36. Ensign, L. M.; Schneider, C.; Suk, J. S.; Cone, R.; Hanes, J., Mucus penetrating nanoparticles: biophysical tool and method of drug and gene delivery. Advanced Materials 2012, 24 (28), 3887–3894.

37. Huckaby, J. T.; Lai, S. K., PEGylation for enhancing nanoparticle diffusion in mucus. Advanced drug delivery reviews 2018, 124, 125–139.

38. Mason, T.; Ganesan, K.; Van Zanten, J.; Wirtz, D.; Kuo, S., Particle tracking microrheology of complex fluids. Physical review letters 1997, 79 (17), 3282.

39. Suk, J. S.; Lai, S. K.; Wang, Y. Y.; Ensign, L. M.; Zeitlin, P. L.; Boyle, M. P.; Hanes, J., The penetration of fresh undiluted sputum expectorated by cystic fibrosis patients by non-adhesive polymer nanoparticles. Biomaterials 2009, 30 (13), 2591–7.

40. Lai, S. K.; O’Hanlon, D. E.; Harrold, S.; Man, S. T.; Wang, Y. Y.; Cone, R.; Hanes, J., Rapid transport of large polymeric nanoparticles in fresh undiluted human mucus. Proc Natl Acad Sci U S A 2007, 104 (5), 1482–7.

41. Seliktar, D., Designing cell-compatible hydrogels for biomedical applications. Science 2012, 336 (6085), 1124–1128.

42. Yu, L.; Ding, J., Injectable hydrogels as unique biomedical materials. Chemical Society Reviews 2008, 37 (8), 1473–1481.

