## Supporting Information for "Designing viscoelastic mucin-based hydrogels"

#### Figure S1

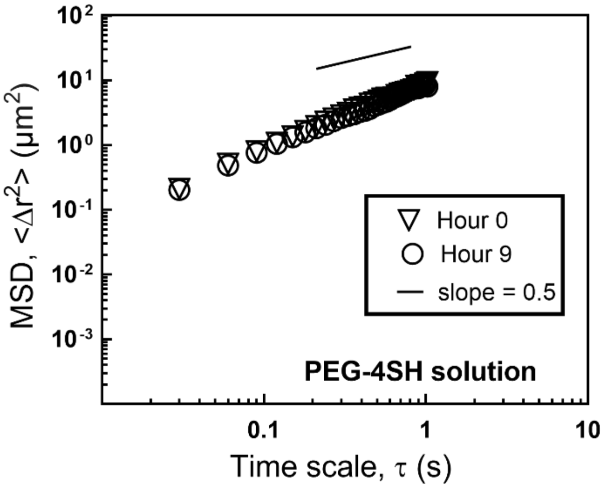


**Figure S1. Microrheology of 2% PEG-4SH solution.** Multiple particle tracking of 100-nm PEG-NP in 2% w/v PEG-4SH polymer solution. The ensemble average mean squared displacement as function of time scale τ (MSD; $\langle \Delta r^2 \rangle$) of 100 nm PEG-NP at hour 0 (triangles) and hour 9 (circles). A reference line with slope *n*=0.5 is included.

### Figure S2

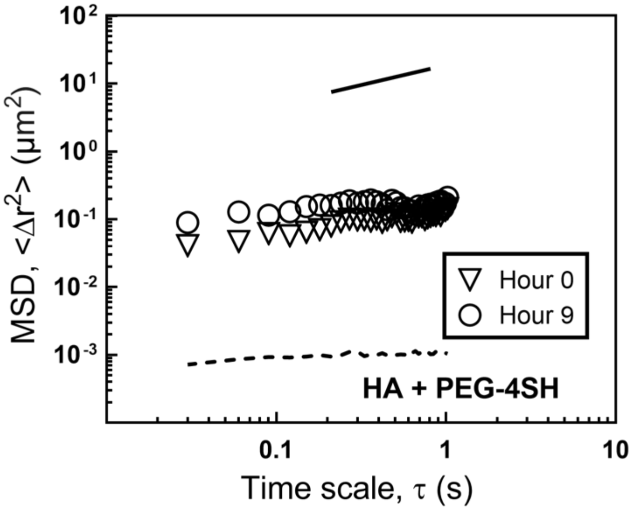


**Figure S2. Microrheology of PEG-4SH mixed with hyaluronic acid.** Multiple particle tracking of 100 nm PEG-NP in 2% w/v PEG-4SH mixed with 2% hyaluronic acid 500 kDa (Lifecore Biomedical). The ensemble average mean squared displacement as function of time scale τ (MSD; $\langle \Delta r^2 \rangle$) of 100 nm PEG-NP at hour 0 (triangles) and hour 9 (circles). Solid line with slope $n$=0.5 and dashed line of 2% PGM/PEG-4SH hydrogel are also shown for reference.

**Figure S3**

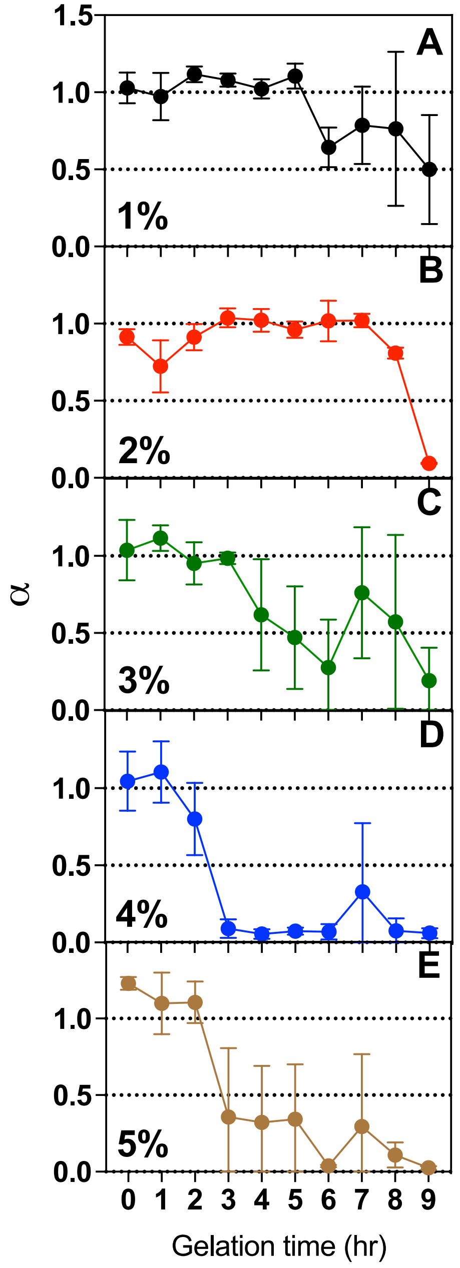


**Figure S3. Gelation rate of mucin-based hydrogels with varying PGM concentration with a constant PEG-4SH concentration of 2% w/v.** Individual experiments using increasing PGM concentrations in w/v of **(A)** 1% (black), **(B)** 2% (red), **(C)** 3% (green), **(D)** 4% (blue), and **(E)** 5% (brown) are shown. Kinetics of PGM gel formation as analyzed by microrheology using 100 nm PEG-NP probes. The gelation point is measured as α= 0.5. Each panel displays the mean and standard error of measured α.

#### Figure S4

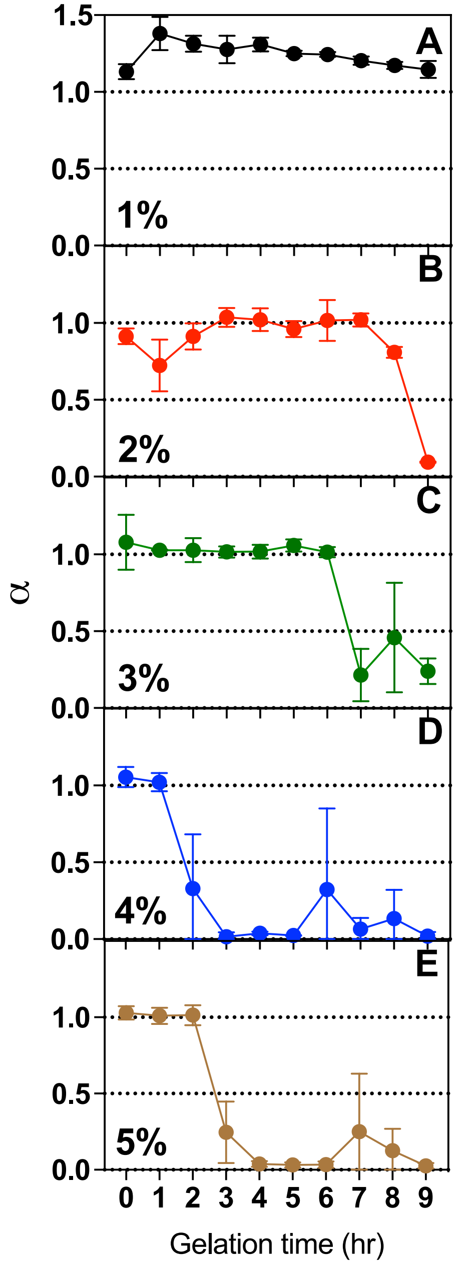


**Figure S4. Gelation rate of mucin-based hydrogels with varying PGM concentration with a constant PEG-4SH concentration of 2% w/v.** Individual experiments using increasing PEG-4SH concentrations in w/v of **(A)** 1% (black), **(B)** 2% (red), **(C)** 3% (green), **(D)** 4% (blue), and **(E)** 5% (brown) are shown. Kinetics of PGM gel formation as analyzed by microrheology using 100 nm PEG-NP probes. The gelation point is measured as α= 0.5. Each panel displays the mean and standard error of measured α.
